# Novel Target Combinations in Lung Squamous Cell Carcinoma proposed by the EMET AI Research Environment and supported by discovery stage experimentation

**DOI:** 10.64898/2026.09.04.749404

**Authors:** Jyothish Soman, Jordan Newington, Simeon M Wong, Nikita Desai, Sophia Leung, Juliana Cudini, Fernando Suarez, Peter Grandsard, Arnab Kundu

## Abstract

Early drug discovery is frequently bottlenecked by target identification, a challenge that becomes particularly difficult in complex diseases driven by overlapping, redundant pathways rather than a single dominant driver. Lung squamous cell carcinoma (LUSC) is one such disease: mutationally complex, lacking a dominant actionable target, and marked by a long series of failed single-agent targeted trials. We hypothesized a research environment, capable of reasoning across interconnected datasets, would be well-suited to generate novel, mechanistically grounded therapeutic hypotheses for LUSC. EMET, an AI research environment utilizing a biomedical knowledge graph of more than 1.5 billion triples connected to over 100 specialised biological databases, was paired with an Agentic Research Director that pursues each hypothesis through a graph-of-thoughts search, invoking scoped link-prediction and retrieval tools and committing every round of findings to persistent memory. The top-ranked hypotheses were two-target combinations rather than a single target. The four highest-ranked combinations were advanced to dose-matrix testing in NCI-H520 and SK-MES-1 cells, and two produced combination effects exceeding either single agent; to our knowledge neither of these two pairings had previously been evaluated in LUSC. Dual inhibition of CDC7 (TAK-931) and PKMYT1 (RP-6306) produced statistically significant synergy in both lines that strengthened from day 5 to day 7 (HSA 19.24 to 24.19 in NCI-H520; 11.07 to 13.73 in SK-MES-1). Dual inhibition of USP13 (spautin-1) and PI3Kα (alpelisib) produced an additive, cell-line-dependent effect, and immunoblotting confirmed the predicted mechanism: time-dependent depletion of MCL-1, c-Myc and SOX-2 driven by the USP13 arm. Agentic reasoning over multi-domain biomedical evidence can therefore nominate testable, mechanism-bearing combination hypotheses that survive experimental scrutiny.

## 1 Introduction

### 1.1 The challenge of novel target discovery

Despite decades of scientific progress, the burden of disease continues to outpace our ability to deliver effective treatments. Thousands of conditions remain without adequate therapy [1], and the traditional drug discovery pipeline is struggling to close that gap: clinical trial failure rates remain above 90% [2], development timelines stretch past a decade [3], and costs now exceed $2 billion per approved drug [4]. Much of this attrition traces back to the earliest stages of discovery, where target selection and hypothesis generation still rely heavily on incomplete biological understanding and trial-and-error experimentation [5, 6]. iterative recycling of failed candidates through early discovery mechanisms amplifies the impact of initial decision-making; thus, marginal gains in early target validation translate into substantial reductions in attrition, development time, and cost (Supplemental Fig 8).

Target discovery is the acute bottleneck of early drug development, presenting the greatest challenge precisely where innovation is needed most. Identifying novel targets requires holistic reasoning across the full complexity of human biology through the integration of multi-omic, clinical, and literature data, moving beyond incremental iterations on validated pathways. However, the sheer breadth of this evidence renders manual analysis intractable, as no single investigator can synthesize connections across millions of publications and hundreds of disparate databases. Inference at scale offers a path forward: systematically traversing biomedical knowledge networks to sharpen target identification, prioritize high-confidence hypotheses, and redirect resources toward candidates most likely to succeed in the clinic.

### 1.2 The state of the art in leveraging AI for novel target identification

Computational target identification and drug repurposing have converged on biomedical knowledge graphs (KGs) encoding genes, proteins, pathways, diseases, and compounds as nodes and their curated or literature-derived relationships as edges. Project Rephetio’s Hetionet with roughly 47,000 nodes and 2.25 million relationships from 29 public resources established that systematic graph-based reasoning could prioritize drug-disease pairs at a scale beyond manual curation [7]. Since then, a growing body of work has applied classical graph algorithms, matrix factorization, and increasingly graph neural networks (GNNs) to KG-based link prediction; recent reviews catalogue dozens of such tools and compare their construction strategies, prediction methods, and limitations [8]. All share a structure: a single query, typically one candidate drug-disease or drug-target pair, is scored against the graph, and top-scoring candidates are returned as hypotheses. In each case the model operates as a global ranking engine. The prediction task is fixed before any biological question is posed and applied uniformly, so a candidate’s visibility depends on its position in one universal ordering rather than its relevance to a specific line of mechanistic reasoning, a distinction we return to in Section 2.2.

A parallel line of work targets combination therapy, framing synergy prediction as supervised learning over large-scale combination screens. DeepSynergy, among the first deep learning approaches, combined chemical and genomic features to predict pairwise synergy scores, but its authors reported a persistent limitation of the paradigm: performance degrades sharply when extrapolating beyond the training distribution of drugs or cell lines [9]. Subsequent GNN methods improved on this baseline by modeling drug structure at multiple scales and disentangling cellline-specific from cell-line-invariant representations [10], yet remain fundamentally data-limited and are as good as the historical screens they are trained on, and therefore poorly suited to nominating untested combinations such as a CDC7 inhibitor with a PKMYT1 inhibitor.

More recently, large language models (LLMs) have been layered onto biomedical KGs, most often via retrieval-augmented generation that lets an LLM query a KG for supporting evidence before answering [11]. This improves interpretability but preserves the single-hypothesis structure of earlier KG methods: the LLM still converges on one answer per query. The EMET platform and the Agentic Research Director used here instead adopt a graph-of-thoughts search, in which an LLM-driven reasoning process represents candidate hypotheses as nodes in an explicit graph, letting independent lines of reasoning branch, be evaluated in parallel, and recombine when they converge on shared mechanistic logic — an approach shown on general-purpose to outperform both single-path chain-of-thought [12] and tree-of-thoughts prompting [13], with substantial gains in solution quality at lower cost [14]. Applied to therapeutic hypothesis generation, this lets the Director reason jointly over a KG and omics data to explore a combinatorial hypothesis space directly, rather than scoring one pre-specified pair at a time, including target combinations with no precedent in screening data. Because graph-of-thoughts search can surface connections that are structurally plausible but poorly evidenced, it is paired with EMET’s validation layer (Section 2.2.5), which cross-references every surviving hypothesis against the literature and experimental evidence base.

#### 1.3 LUSC as a candidate for AI-driven target discovery

Lung cancer is the most commonly diagnosed malignancy and the leading cause of cancer death worldwide, with an estimated 2.5 million new cases and 1.8 million deaths in 2022 [15]. Lung squamous cell carcinoma (LUSC), the second most common histological subtype after adenocarcinoma, accounts for roughly 25–30% of non-small cell lung cancer and nearly 30% of lung cancers diagnosed in men globally [16]. Despite progress in other subtypes, advanced LUSC carries five-year survival below 10% in metastatic disease and median overall survival of only 10–12 months on standard platinum-based chemotherapy [17].

This prognosis is rooted in LUSC’s molecular architecture. The Cancer Genome Atlas identified LUSC as one of the most mutationally complex solid tumors, with near-universal inactivation of TP53 (>90%) and CDKN2A alongside recurrent amplification of SOX2, PIK3CA, and FGFR1 [18]. Unlike EGFR- or ALK-driven adenocarcinoma, LUSC lacks a single dominant druggable driver; its pathogenesis reflects coordinated, redundant dysregulation across interacting pathways [19]. The recurrent alterations that do exist are predominantly high-frequency somatic mutations in markers rather than activating mutations in a target protein; lacking causal genetic association evidence, they define a complex, lineage-specific tumor state rather than an actionable node whose isolated inhibition collapses the disease.

That distinction explains why strategies that transformed lung adenocarcinoma (LUAD) have not transferred. In LUAD, identifying and inhibiting the driver produces large, durable responses; in LUSC the same playbook has repeatedly failed, even when carefully executed. FGFR inhibition with molecules such as AZD4547 and Rogaratinib being the clearest examples [20–22]. Anti-angiogenics were closed off entirely in squamous histology after fatal pulmonary hemorrhage [23], and PI3K inhibitors have repeatedly failed as monotherapy despite frequent PIK3CA amplification [24]. Immunotherapy and chemotherapy therefore remain the systemic backbone, alongside surgery and radiation for localized disease, leaving a pronounced unmet need for more effective targeted strategies in this histology [24].

LUSC is consequently an unusually informative test case for AI-driven target discovery. A disease whose biology is redundant and distributed, whose actionable alterations are numerous but individually insufficient, and whose single-target trials have a documented record of failure is precisely where hypothesis-by-hypothesis manual search performs worst — and where reasoning across the whole evidence base at once has most to add.

Our contribution is threefold. First, we show how the director surfaced two novel combinatorial hypotheses for LUSC — dual inhibition of CDC7 and PKMYT1, and of USP13 and PI3K — by integrating multi-omic data, functional and pathway-level evidence, pharmacological and drug safety data, and clinical/translational and literature-derived knowledge (Section 2.2). Second, we lay out the mechanistic rationale supporting and contextualizing each (Sections 3.4–3.5). Third, we report preclinical dose-matrix experiments testing both and their results (Sections 3–4).

## 2 Methods

### 2.1 The EMET platform design

We have deployed EMET, an AI Research Environment, built by BenchSci for this study. Architecturally, the system comprised a research environment (EMET) and a specialized capability for novel LUSC hypothesis generation (Agentic Research Director) described in Section 2.2. The Agentic Research Director is an instance of a specialized capability that can be created in the EMET environment. Figure 1 illustrates the architecture of the EMET platform and the Agentic Research Director

**Figure 1.**
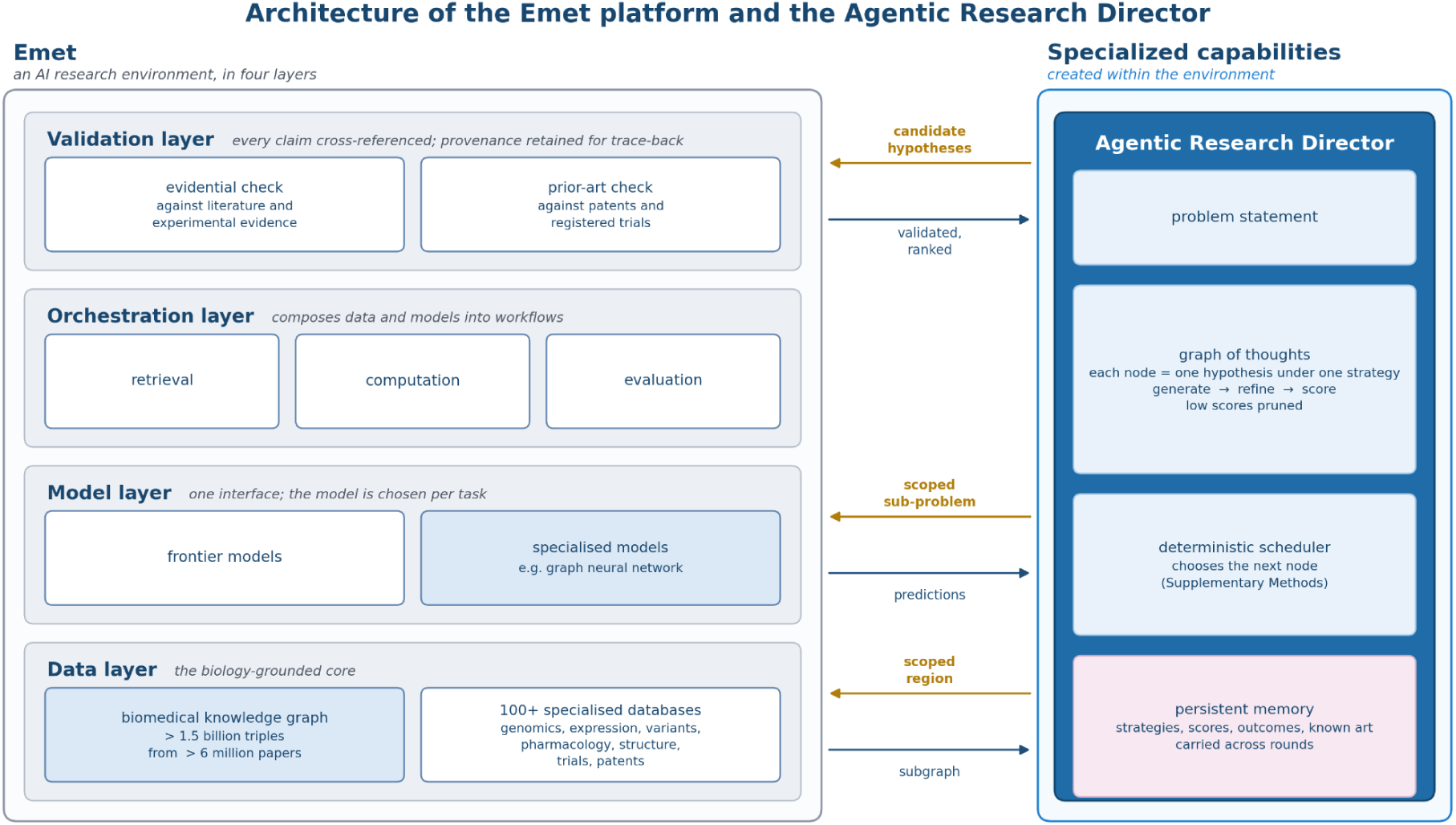
Overall architecture of the EMET platform and the specialized Agentic Research Director. The Agentic Research Director relies on the EMET platform for accessing data and models while performing its evaluations which are then validated through EMET’s Validation layer.

**Figure 2.**
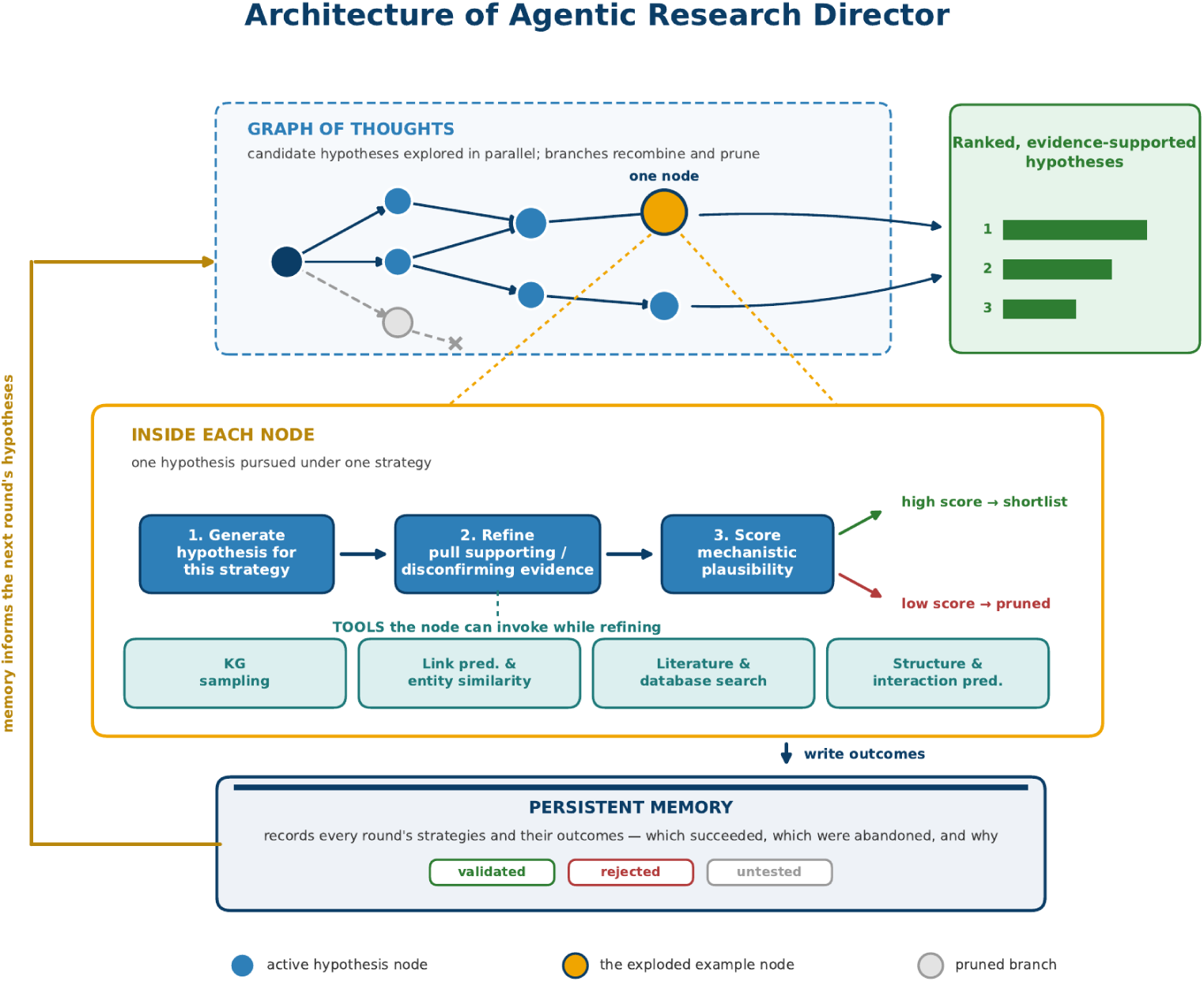
Architecture of Agentic Research Director. The Director maintains a graph of thoughts in which each node is one hypothesis pursued under one strategy. Every node runs the same internal generate– refine–score cycle and, during refinement, can invoke a set of machine learning and retrieval tools, such as knowledge-graph sampling, link-prediction and entity-similarity models, literature and database search, and structure- and interaction-prediction models, scoped to the region its strategy makes relevant. High- scoring nodes are carried forward to the platform’s validation and prioritisation layer, while low-scoring nodes are pruned. A persistent memory records each round’s strategies and their outcomes, and feeds that history into the next round.

EMET provides the environment linking biomedical data, models, bioinformatics tools, specialized workflow-centric agents, specialized point solutions and lab-in-the-loop automation capabilities. EMET is designed to address a fundamental structural problem in the drug discovery stack. The scientific knowledge and tools required to support evidence-based research workflows, novel target discovery being one, is currently distributed across hundreds of sources and toolkits. To solve this, the EMET platform is designed in 4 layers - a Data Layer, a Model Layer, an Orchestration Layer and a Validation Layer

At the core of the *Data Layer* is a biomedical knowledge graph spanning more than 1.5 billion triples, with relationships derived from over 6 million papers, supporting canonicalization of genes, diseases, drugs, and other biomedical terms to a common identity system, alongside relationship, variant, and evidence enrichment across distinct biomedical predicates.

This knowledge graph is in turn connected to more than 100 specialized biological databases spanning the major domains summarized in Table 1. It is this breadth and depth of interconnected evidence — rather than any single database or data type — that allows the Agentic Research Director (described in Section 2.2) to reason jointly across mechanism, genomics, pharmacology, and clinical outcomes when generating and validating combinatorial hypotheses.

**Table 1.** Data domains underlying EMET’s knowledge graph, connecting more than 100 specialized biological databases.

| Domain | Representative Sources |
| --- | --- |
| Literature and evidence | PubMed, Europe PMC, OpenAlex |
| Cancer genomics | cBioPortal, TCGA, DepMap |
| Gene expression | GTEx, single-cell atlases |
| Clinical and variant data | ClinVar, gnomAD |
| Drug discovery and pharmacology | ChEMBL, Open Targets |
| Protein structure | AlphaFold, PDB |
| Pathway and interaction networks | Reactome, STRING |
| Spatial and anatomical atlases | Reference anatomical atlases |
| Clinical trials | US clinical trials |
| Patents | USPTO |

A *model layer* exposes both frontier and specialised models behind a uniform interface, with the right model is selected for each scientific task. For instance, a custom graph neural network was used for scoped link prediction (elaborated in Section 2.2.2).

An *orchestration layer* composes these into workflows, each a defined sequence of retrieval, computation and evaluation steps addressed to a recognizable research question.

Finally, a *validation layer* cross-references any claim against the underlying literature and experimental evidence base, retaining the provenance so that an output can be traced back to the records supporting it; this is the layer that assesses candidate hypotheses in Section 2.2.5.

### 2.2 The Agentic Research Director

The Agentic Research Director is a specialized capability within the EMET platform, distinguished from the computational approaches reviewed in Section 1.2 by one property: it runs multiple computational and reasoning experiments, learns from them, and returns a consolidated outcome. Given a problem statement, it pursues multiple hypotheses in parallel, validating each with data, machine learning, and memory tools, then uses what it learns to define the next frontier from the system’s most current information — much as a human research director coordinates a team of scientists, each working a different candidate.

Candidates come from a graph-of-thoughts search in the sense of Besta and colleagues, where the units of reasoning a language model produces are vertices and their dependencies are edges, so independent lines of reasoning can branch, be evaluated in parallel, and recombine [14]. Each node is one hypothesis under one strategy, and every node runs the same cycle: generate a candidate consistent with its strategy, refine it against supporting or disconfirming evidence, and score it for mechanistic plausibility and evidential support, pruning low scorers rather than carrying them forward. The search spans two spaces at once. Strategy space is the set of reasoning angles available — synthetic-lethal dependency, protein-interaction convergence, expression-based vulnerability, precedent from validated combinations in other indications — and options space is the concrete genes, drugs, and pathways within each. Rather than fixing a strategy and enumerating its options, the Director navigates both together, so a promising strategy opens new options and a promising option motivates a new strategy. Node scheduling is decided by a deterministic outer loop rather than model discretion; that loop, and the division of labour between model- and code-driven steps, is described in the Supplementary Methods.

A second differentiator is that conventional knowledge-graph pipelines run a predictive model once over the entire space (every gene–disease or drug pair) to produce a single context-free ranking, with the prediction task fixed before any biological question is posed, so a candidate’s visibility depends on its position in one universal ordering. The Director instead operates the same class of model as a context-specific instrument whose task is constructed per line of reasoning. That difference makes candidates reachable that a global ranking would bury, and everything below exists to support it.

Scoped model use is where that distinction becomes concrete. A node first samples the knowledge graph in the region its strategy makes relevant, then specifies what to predict over that region: which entities, which relationship type, in which direction. That scoped sub-problem, not the whole graph, is passed to the graph neural network. Predictions are read back and interpreted relative to the requesting strategy, and may motivate a narrower follow-up query, so several models compose within one line of reasoning, each output conditioning the next input. One model therefore yields different, locally relevant answers depending on which branch is asking, because both input scope and interpretive frame change with the branch. The CDC7/PKMYT1 pairing illustrates the consequence. With no precedent in any combination screen, a model ranking drug pairs by expected synergy across the whole space has no signal to place it highly, whereas the narrower question of which kinase dependencies become essential when origin firing is suppressed in cells whose G1 checkpoint is already dismantled meets locally dense evidence, and the pairing is reachable.

Beyond a single search, the Director pursues a research mission across multiple rounds rather than returning a one-shot answer. Each round’s outcomes, strategies attempted, scores achieved, and candidates retained, rejected for want of support, or set aside as already claimed, are committed to a persistent memory outliving the round, so later rounds extend what worked, deprioritize what did not, and avoid re-deriving settled conclusions.

### 2.3 In vitro validation of hypotheses

All in vitro experiments reported here were performed under contract by Pharmaron (Beijing, China) to protocols agreed with the authors..

#### 2.3.1 Selection of cell lines and tool molecules

To test both hypotheses, two well-characterized human LUSC models were selected: NCI-H520 (derived from a primary squamous cell carcinoma) and SK-MES-1 (derived from a pleural metastasis). These widely established cell lines were chosen to represent distinct clinical stages of LUSC.

For the CDC7/PKMYT1 hypothesis, both targets have clinical-stage inhibitors with well-characterized pharmacology: TAK-931 for CDC7 [25, 26] and RP-6306 (lunresertib) for PKMYT1 [27, 28]. For the USP13/PI3K hypothesis, only the PI3K arm has a clinical-stage agent (alpelisib). No selective USP13 inhibitor is commercially available. Hence USP13 was targeted pharmacologically with spautin-1, which is known to co-inhibit USP10, and genetically by siRNA-mediated knockdown for greater selectivity.

#### 2.3.2 Cell culture and dosing

For the CDC7/PKMYT1 assay, NCI-H520 and SK-MES-1 cells were seeded in 384-well plates in 40 μL of complete growth medium at densities optimized for each line (NCI-H520: 3,200 cells/well; SK-MES-1: 1,200 cells/well) and allowed to adhere prior to compound addition. TAK-931 (MCE, HY100888) and RP-6306 (MCE, HY-145817A) were tested in a 7×7 dose matrix at 0, 1, 10, 30, 100, 300, and 1000 nM each, alone and in all pairwise combinations, with plates incubated at 37^∘^C, 5% CO_2_ for 5 d or 7 d post-treatment, with four independent replate plates per cell line and time point.

For the USP13/PI3K assay, the same two lines were seeded in 96-well plates in 100 μL of complete growth medium at densities optimized for each line (NCI-H520: 10,000 cells/well; SK-MES-1: 4,000 cells/well) and incubated overnight at 37^∘^C, 5% CO_2_ prior to compound addition. Alpelisib (MCE, HY-15244) and spautin-1 (MCE, HY-12990) were tested in a 5×7 dose matrix (spautin-1: 0, 1, 5, 10, and 20 μM; alpelisib: 0, 1, 5, 10, 15, 20, and 30 μM), alone and in all pairwise combinations, with plates incubated at 37^∘^C, 5% CO_2_ for 48 h post-treatment. This assay was run as four independent replicate plates per cell line.

#### 2.3.3 Cell viability assay

At each timepoint, CellTiter-Glo 2.0 Reagent was added to assay plates (20 μL for the 384-well CDC7/PKMYT1 assay; 50 μL for the 96-well USP13/PI3K assay), plates were centrifuged (1 min, 1000 rpm), and luminescence, proportional to cellular ATP content, was recorded after a 20-minute incubation using an EnSight multimode plate reader. Viability was therefore assessed by a single bulk endpoint in both experiments.

#### 2.3.4 siRNA-mediated USP13 knockdown and treatment groups

To evaluate USP13-specific effects versus the dual USP10/USP13 inhibitor spautin-1 alongside PI3K inhibition, NCI-H520 (3×10^5^ cells/well) and SK-MES-1 (3.5×10^5^ cells/well) cells were seeded in 12-well plates in 1 mL complete medium and cultured overnight (37^∘^C, 5% CO_2_). Cells were transfected with 50 nM USP13 siRNA (sense: 5′-GAUCAUUGUUCACAUGGAAdTdT-3′; antisense: 5′-UUCCAUGUGAACAAUGAUCdTdT-3′) or a scrambled control (GenScript) using Lipofectamine RNAiMAX in Opti-MEM (100 µL complex/well). Transfection was maintained for 48 h before downstream treatment.

After the 48 h knockdown period, cells were reseeded into 12-well plates and assigned to one of the following treatment groups: (i) vehicle control (DMSO); (ii) non-targeting control (50 nM scrambled siRNA); (iii) USP13 knockdown (50 nM USP13 siRNA); (iv) USP13 inhibition (10 μM spautin-1); (v) PI3Kα inhibition (5 μM alpelisib); (vi) genetic–pharmacological combination (50 nM USP13 siRNA, 48 h pre-treatment, followed by 5 μM alpelisib); and (vii) pharmacological combination (10 μM spautin-1 + 5 μM alpelisib). Lysates were collected at 1, 4, 24, 48 and 72 h after treatment, of which the 24, 48 and 72 h timepoints are reported in Section 3.6.

#### 2.3.5 Western blot analysis

Cells were lysed in pre-chilled RIPA buffer with protease and phosphatase inhibitors (30 min, 4°C). Lysates were clarified by centrifugation, quantified by BCA assay, diluted in 4× SDS loading buffer, denatured (70°C, 10 min), and stored at −80°C.

Equal protein (15 µg/lane) was resolved on 4–12% Bis-Tris SDS-PAGE (80 V, 30 min; then 120 V, 1.5 h) and transferred to PVDF (300 mA, 60 min, 20% methanol). Membranes were blocked in 5% BSA/TBST (1 h, RT), probed with primary antibody overnight at 4°C, then with HRP-conjugated secondary (1 h, RT). After TBST washes (3 × 7 min per step), bands were visualized by enhanced chemiluminescence or Odyssey imaging. Primary antibodies and working dilutions are listed in Table 2; β-actin served as loading control.

**Table 2.** Primary antibodies used for immunoblot analysis of the USP13/PI3K arm. β-actin served as the loading control.

| Target | Supplier, catalog no. | Dilution |
| --- | --- | --- |
| MCL-1 (rabbit mono-clonal) | Cell Signaling Technology, 5453 | 1:1000 |
| USP13 | Abcam, ab109264 | 1:1000 |
| PIK3CA | Abcam, ab40776 | 1:1000 |
| SOX2 | Abcam, ab97959 | 1:1000 |
| c-Myc | Abcam, ab32072 | 1:1000 |
| β-actin (8H10D10, mouse mAb) | Cell Signaling Technology, 3700 | 1:10000 |

Band intensities were quantified in Image Studio Lite (LI-COR Biosciences), with each target band normalized to β-actin on the same blot and relative expression reported as that ratio.

#### 2.3.6 Synergy analysis

Percent inhibition relative to vehicle-treated wells was calculated for each dose-matrix well and analyzed in SynergyFinder+ (https://synergyfinder.org, version [R-3.10.3], accessed [2026-08-14]) [29]. Combinations were scored under the Highest Single Agent (HSA) model [30], which sets the expected effect as the larger of the two single-agent effects at the corresponding concentrations; selected combinations were also scored under Bliss independence [31], which treats the agents as acting through probabilistically independent processes. The synergy score is the mean excess of observed over expected effect across the matrix. P values accompanying whole-matrix scores are those returned by SynergyFinder+, testing mean synergy against a null of no interaction. Because HSA and Bliss rest on different nulls, their scores are not directly comparable and are reported separately by model throughout [32]. By the convention used here, scores below −10 indicate antagonism, −10 to 10 additivity, and above 10 synergy.

## 3 Results

### 3.1 Combination therapies emerge as the leading therapeutic strategy

Asked to identify novel therapeutic targets in LUSC without a pre-specified target list or drug class, the EMET platform with the Agentic Research Director returned dozens of novel, evidence-validated combinatorial therapeutic hypotheses spanning distinct mechanistic strategies, from which the 20 top-ranked mechanisms with the strongest supporting evidence were carried forward. Each surviving candidate arrived with an explicit mechanistic rationale and its supporting provenance attached, having passed both the Director’s internal plausibility scoring and the EMET’s independent validation layer that deprioritizes hypotheses lacking adequate support in the literature and experimental evidence base, or are not novel.

Following the ranking approach outlined above, all top-ranked hypotheses were combination therapies, even though the system was not explicitly prompted to seek multi-target strategies. This is not unexpected, and likely reflects the fact that LUSC is characterized by multiple driver alterations rather than a single dominant one.

### 3.2 Top Four hypotheses prioritized for in vitro validation

The top four hypotheses were selected for *in vitro* validation (Table 3). The four differ meaningfully in translational maturity: some pair two targets that already have clinical-stage inhibitors, while others pair a well-precedented target with one that remains at an earlier, preclinical stage of drug discovery.

**Table 3.** The four combinatorial hypotheses selected for in vitro validation, following internal vetting of the 20 top-ranked EMET-generated candidates for LUSC.

| Targets | Why Novel for LUSC | Translational Maturity |
| --- | --- | --- |
| CDC7 + PKMYT1 | Predicted “double-jeopardy” mitotic catastrophe: CDC7 inhibition generates under-replicated DNA while PKMYT1 inhibition removes the checkpoint that would otherwise allow its repair before mitosis | Both targets clinical-stage (TAK-931, RP-6306); combination untested |
| USP13 + PI3K | Exploits the 3q26 co-amplification of USP13, SOX2, and PIK3CA; USP13 inhibition is predicted to create a synthetic vulnerability that reopens a therapeutic window for PI3K inhibitors | PI3K clinical-stage; USP13 preclinical only; combination tested in other solid tumors [33] |
| POLQ + TERT | TERT inhibition generates telomeric double-strand breaks that LUSC cells resolve via POLQ-mediated end joining; blocking POLQ is predicted to convert a slow-acting telomerase inhibitor into an acute source of irresolvable genomic damage. | Both targets clinical-stage (ART4215, Imetelstat); combination untested |
| GOLPH3 + TERT | Pairs an acute, mTOR-linked survival-signaling node with the long-term replicative-immortality node on a shared chromosome 5p amplicon (5p13 / 5p15.33); GOLPH3's role in mTOR signaling is mechanistically linked to TERT transcription. | No GOLPH3 inhibitor yet in development; highest-risk combination of the four |

In in vitro testing, two hypotheses produced measurable combination effects exceeding single-agent treatment in early preclinical assays: dual inhibition of CDC7 + PKMYT1 and of USP13 + PI3Kα (full results in Sections 3.4, 3.5, 3.6). Neither of these two combinations had, to our knowledge, been tested together in a LUSC context before this study, and both are characterized in detail below.

The remaining two combinations were less conclusive. ART558 (POLQ inhibitor, 10 µM) with BIBR1532 (TERT inhibitor, 25 µM) gave moderate growth inhibition at 96 h (data not shown). This is mechanistically unsurprising: telomerase inhibition kills via progressive telomere attrition, and POLQ end-joining substrate accumulates only over successive divisions, so a 96 h end-point may sample the interaction before damage accrues. Extended-timepoint dose matrices are recommended to test for a larger effect. For GOLPH3 + TERT, no commercial GOLPH3 inhibitor exists, so GOLPH3 was siRNA-depleted alongside BIBR1532 (25 µM); no consistent viability trend emerged across cell lines and timepoints (data not shown).

### 3.3 CDC7/PKMYT1 co-inhibition produces synergistic growth inhibition

The first hypothesis exploits the coupling between replication licensing and mitotic entry. CDC7, as the catalytic subunit of the DBF4-dependent kinase, phosphorylates the MCM2–7 helicase to license and fire replication origins, while PKMYT1 is the dominant kinase phosphorylating CDK1 at Thr14, holding the CDK1–cyclin B complex inactive through S phase and G2. Inhibiting CDC7 leaves late origins unfired and replication incomplete; inhibiting PKMYT1 removes the brake that would otherwise arrest those same cells before division. Cells are therefore driven into mitosis carrying under-replicated chromosomes, a ‘double-jeopardy’ convergence on two non-overlapping checkpoints predicted to terminate in mitotic catastrophe. LUSC is an unusually permissive setting for this strategy because near-universal TP53 loss has already dismantled the G1 checkpoint, leaving the G2/M transition as the only remaining safeguard [18, 34], and both arms are independently druggable with clinical-stage inhibitors [25–28].

Combining TAK-931 and RP-6306 produced a synergistic dose-dependent growth inhibition in both LUSC lines that exceeded either single agent alone across the tested concentration range, with the interaction strengthening from day 5 to day 7 post-treatment (Figure 3A and B). In NCI-H520 cells, mean dose-matrix inhibition was 26.42% at day 5 and 30.26% at day 7, and the overall HSA synergy score was 19.24 (p=1.15×10^−5^) at day 5, increasing to 24.19 (p=5.45×10^−6^) at day 7 (Figure 3A). In SK-MES-1 cells, mean dose-matrix inhibition was 32.42% at day 5 and 39.26% at day 7, with an overall HSA synergy score of 11.07 (*p* = 1.89 × 10^−4^) at day 5 and 13.73 (p=4.76×10^−5^) at day 7 (Figure 3B). By the thresholds applied here the whole-matrix interaction classifies as synergistic in both cell lines and at both timepoints, with the synergy score highly statistically significant in every condition (p<2×10^−4^ in all four). The same conclusion holds under an alternative reference model. Recomputing synergy over the identical dose matrices under Bliss independence returned concordant scores that likewise increased from day 5 to day 7 (Supplementary Figure 2). In NCI-H520 cells the Bliss synergy score was 13.49 at day 5 (p = 6.78 × 10^−101^) and 21.65 at day 7 (p = 9.70 × 10^−65^), placing the interaction in the synergistic range at both timepoints. In SK-MES-1 cells the Bliss score was 6.48 at day 5 (p = 1.17 × 10^−56^) and 12.10 at day 7 (p = 4.94 × 10^−39^), so this line falls in the upper additive range at day 5 — below the HSA estimate of 11.07 — and crosses into the synergistic range by day 7. The direction of the interaction and its strengthening with time are therefore independent of the reference model chosen.

**Figure 3.**
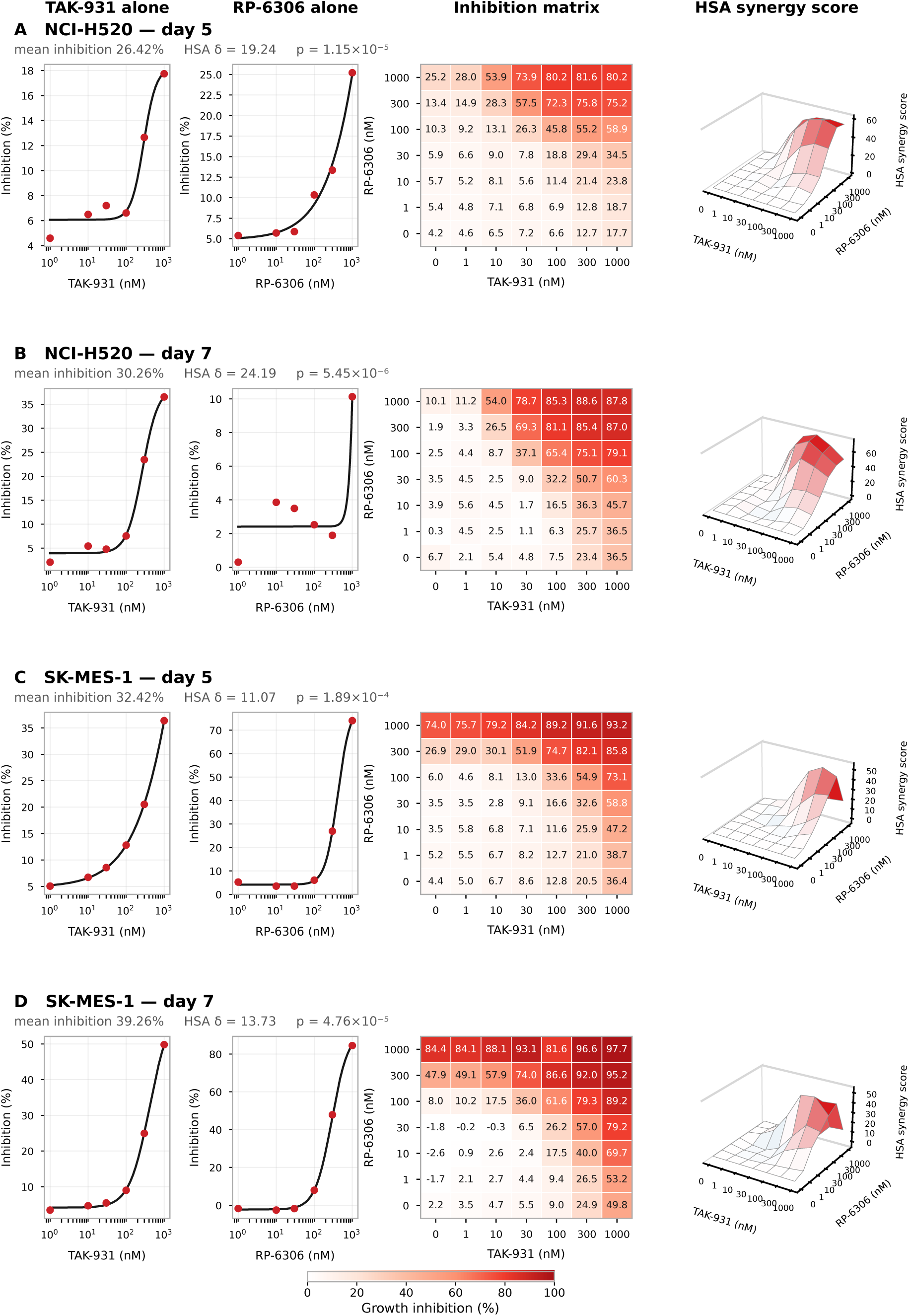
Synergistic and time-dependent growth inhibition induced by dual CDC7 and PKMYT1 inhibition in LUSC cell lines. For each cell line and timepoint, from left to right: single-agent dose-response curves for TAK-931 and for RP-6306, the replicate-averaged dose-response inhibition matrix, and the three-dimensional HSA synergy score surface. (A) NCI-H520 at day 5. (B) NCI-H520 at day 7. (C) SK-MES-1 at day 5. (D) SK-MES-1 at day 7. The whole-matrix mean HSA synergy score (δ) and its p-value are indicated above each row of panels. Both agents were tested at 0, 1, 10, 30, 100, 300 and 1000 nM in a 7 × 7 dose matrix over a 384-well CellTiter-Glo 2.0 viability assay; synergy scores above 10 indicate a synergistic interaction. All panels were analysed in SynergyFinder+ using the HSA reference model [29, 30].

### 3.4 USP13 co-inhibition widens the therapeutic window for PI3K inhibition in NCI-H520 cells

The second hypothesis targets two non-overlapping regulatory layers that converge on the same survival effectors. USP13 is among the most frequently amplified genes in LUSC, sitting on the 3q26 amplicon alongside PIK3CA and SOX2, and acts post-translationally as a deubiquitinase that stabilizes c-MYC and reinforces the squamous lineage program [35]; in other epithelial cancers it likewise stabilizes the anti-apoptotic protein MCL-1 [36]. PI3K/AKT/mTOR signaling, constitutively active in this genomic context, independently drives the de novo production of the same effectors. Because one arm operates on protein stability and the other on signal transduction, resistance to either is unlikely to confer cross-resistance, and co-inhibition is predicted to deplete the existing pool of pro-survival proteins while simultaneously blocking its replenishment, lowering the apoptotic threshold below what either agent achieves alone. Critically, the therapeutic aim here is not supra-additive killing in its own right but the reopening of a therapeutic window for PI3K inhibitors, a class that has repeatedly failed as LUSC monotherapy [24], by neutralizing USP13-dependent survival signaling that has been hypothesized to confer resistance to PI3K inhibitors in ovarian cancer [33].

Combining alpelisib and spautin-1 produced an additive dose-dependent growth inhibition in both LUSC lines at 48 h (Figure 4A and B). In NCI-H520 cells, mean dose-matrix inhibition was 36.91% (±1.25% SD across replicates), with an overall HSA synergy score of 7.86 (Figure 4A). In SK-MES-1 cells, mean dose-matrix inhibition was 39.68% (±0.97% SD), with an overall HSA synergy score of 1.58 (Figure 4B). By the thresholds applied here both remain within the additive range. Scoring the same replicate-averaged matrices under the Bliss independence reference model gives a more conservative reading of the interaction (Supplementary Figure 3).

**Figure 4.**
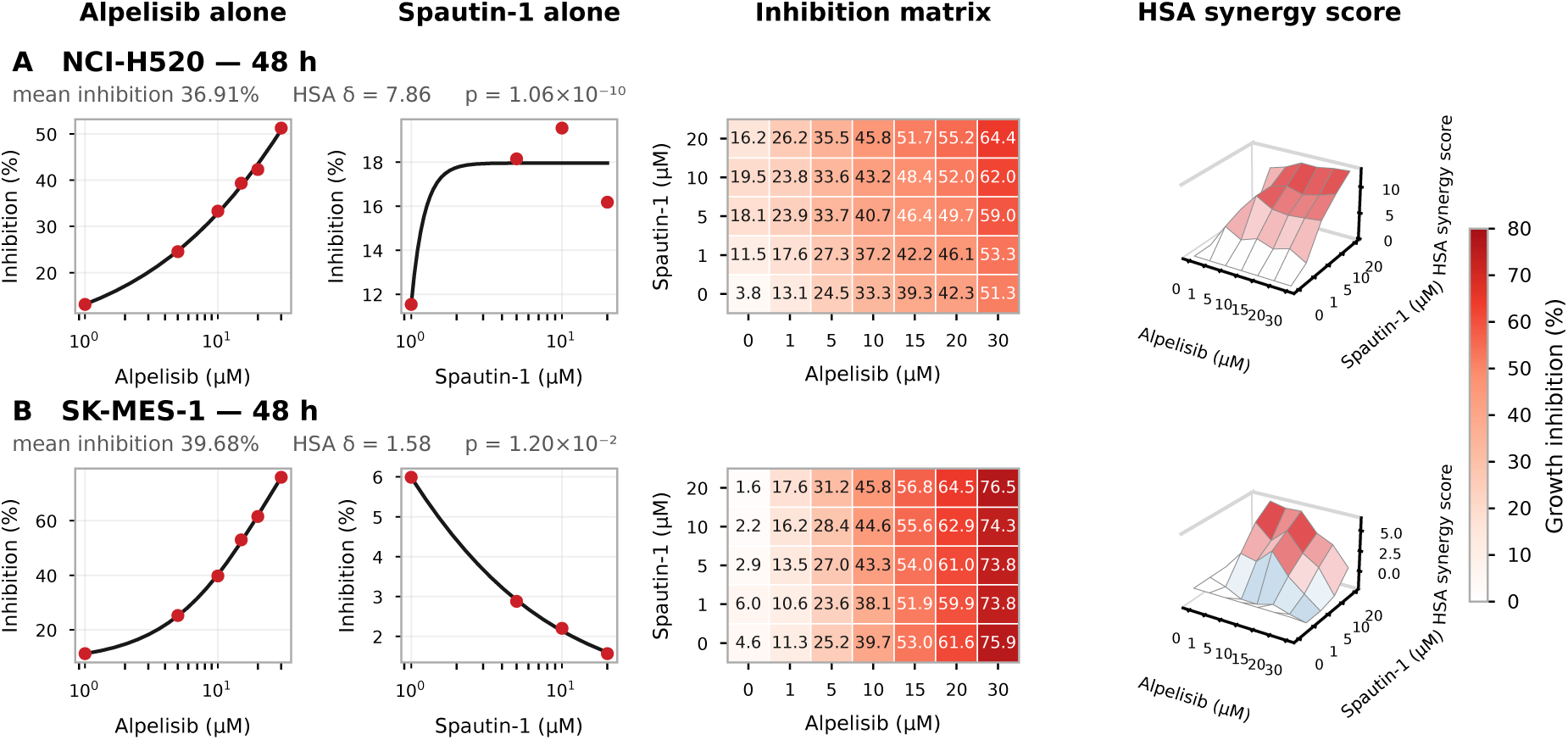
Combined alpelisib and spautin-1 treatment yields cell-line-specific additive growth inhibition in LUSC cells. For each cell line, from left to right: single-agent dose-response curves for alpelisib and for spautin-1, the replicate-averaged dose-response inhibition matrix, and the three-dimensional HSA synergy score surface, all at 48 h. (A) NCI-H520 (B) SK-MES-1. The whole-matrix mean HSA synergy score (δ) and its p-value are repeated above each row of panels. Spautin-1 was tested at 0, 1, 5, 10 and 20 µM and alpelisib at 0, 1, 5, 10, 15, 20 and 30 µM in a 5 × 7 dose matrix over a 96-well CellTiter-Glo 2.0 viability assay, with four independent replicate plates per cell line; both panels show the replicate-averaged matrices. All panels were analysed in SynergyFinder+ using the HSA reference model [29, 30].

The inhibition matrices (Figure 4 A, B) demonstrate that, even though spautin-1 in isolation does not show an appreciable dose response, the presence of spautin-1 increases the inhibition potential of alpesilib. For instance, in NCI-H520 cells, 30 μM of alpesilib achieves ∼51% inhibition in isolation while a very comparable degree of inhibition is achieved with 15 μM of Alpesilib in the presence of 20 μM of spautin-1. A similar but quantitatively smaller effect is observed in the inhibition matrix in SK-MES-1 cells. This supports the hypothesis that co-inhibition of USP13 increases the therapeutic window of PI3Kα inhibitors.

### 3.5 PI3Kα Inhibition Enhances USP13 Target Depletion in LUSC cells

The viability data establish that the two arms interact, but not that they act through the predicted mechanism. The hypothesis makes a specific molecular prediction: if USP13 sustains LUSC by post-translationally stabilizing MCL-1, c-Myc and the squamous lineage factor SOX-2, then removing USP13 function should lower the steady-state level of those proteins, and blocking their PI3K-driven replenishment with alpelisib should deepen and extend the loss. We therefore assayed the same two cell lines by immunoblot at 24, 48 and 72 h across a seven-condition matrix combining USP13 depletion (siRNA) or inhibition (10 μM spautin-1) with the PI3Kα-selective inhibitor alpelisib (5 μM) (Figure 5). USP13 siRNA reduced USP13 protein to 12% and 7% of the non-targeting control at 24 and 48 h respectively, confirming target engagement in the depletion arm (Supplementary Figure 2).

**Figure 5.**
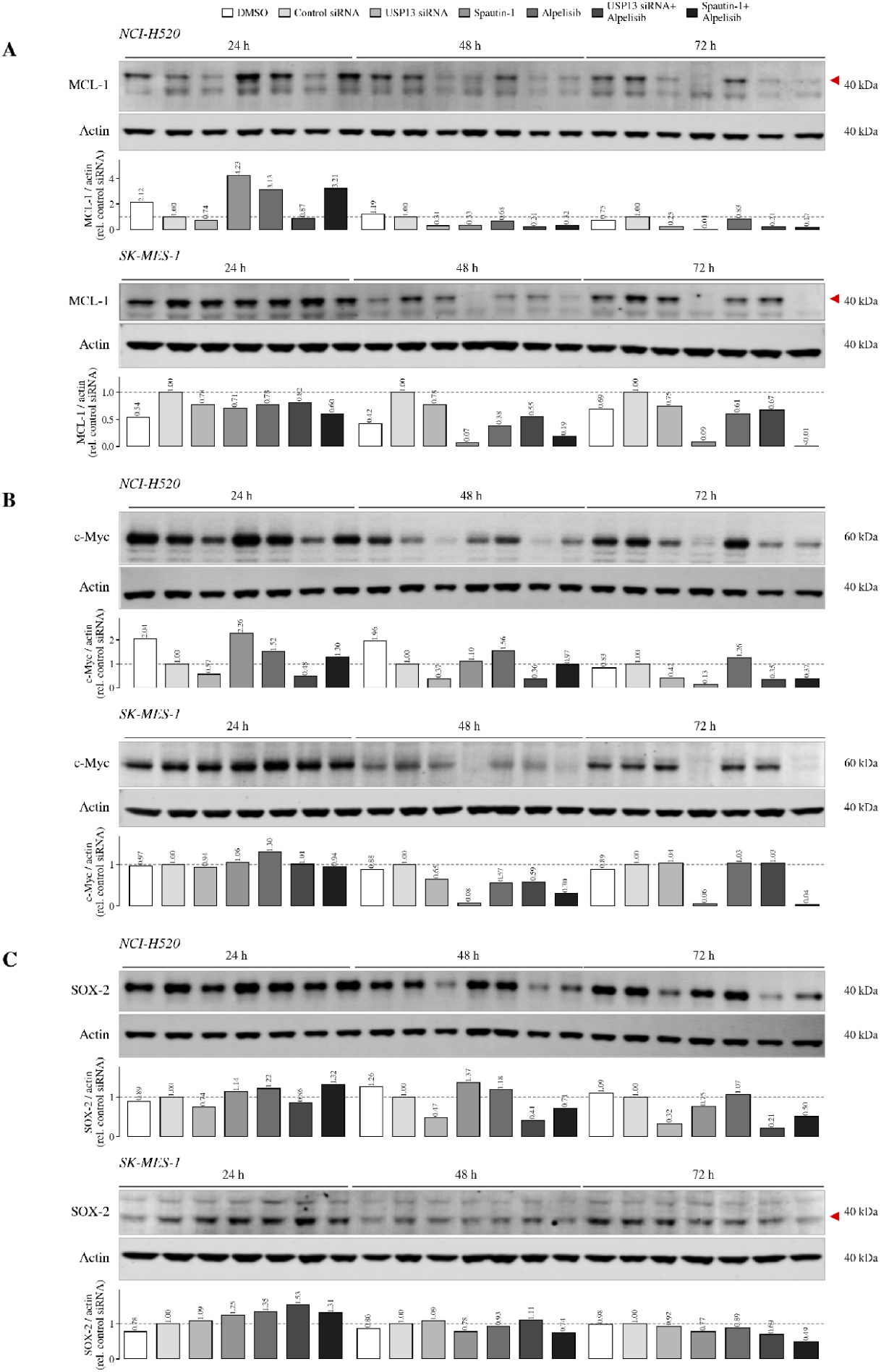
MCL-1, c-Myc and SOX-2 protein levels following USP13 loss of function and PI3K inhibition in lung squamous cell carcinoma lines. Immunoblot analysis of the LUSC lines NCI-H520 and SK-MES-1 at 24, 48 and 72 h following USP13 depletion or inhibition, alone and in combination with the PI3K-selective inhibitor alpelisib. Within each time block the seven lanes are, in order: (1) vehicle (DMSO); (2) non-targeting control siRNA; (3) USP13 siRNA; (4) 10 µM spautin-1; (5) 5 µM alpelisib; (6) USP13 siRNA + 5 µM alpelisib; (7) 10 µM spautin-1 + 5 µM alpelisib. Treatments are identified by the key above each blot. (A) MCL-1 (40 kDa) in NCI-H520 (upper) and SK-MES-1 (lower); red arrowheads indicate the MCL-1 band. (B) c-Myc (60 kDa) in NCI-H520 (upper) and SK-MES-1 (lower). (C) SOX-2 (40 kDa) in NCI-H520 (upper) and SK-MES-1 (lower); the red arrowhead marks the SOX-2 band in SK-MES-1. Actin was probed on the corresponding membrane as the loading control for every panel, and approximate molecular weights are indicated to the right of each blot. Bar graphs beneath each panel show densitometric quantification normalized to actin and expressed relative to the non-targeting siRNA control (dashed line = 1.00). Primary antibodies are as listed in Table 2. Target engagement for USP13 knockdown and spautin-1 treatment, together with PIK3CA protein levels across the same treatment matrix, is shown in Supplementary Figure 4.

The predicted direction was observed, and it was time-dependent. In NCI-H520 cells, densitometry normalized to actin and expressed relative to the non-targeting control siRNA lane within each time block showed no MCL-1 loss at 24 h; by 48 h USP13 siRNA had reduced MCL-1 to 0.31 of control and spautin-1 to 0.33, and by 72 h to 0.25 and 0.01 respectively. Alpelisib alone was comparatively inert at this node (0.68 at 48 h, 0.83 at 72 h), consistent with MCL-1 in this context being maintained primarily by protein stabilization rather than by de novo synthesis downstream of PI3Kα. The lowest MCL-1 levels in the matrix occurred in the combination lanes at both later timepoints (USP13 siRNA + alpelisib, 0.21 at 48 h and 0.21 at 72 h; spautin-1 + alpelisib, 0.32 and 0.17). SOX-2 followed the same pattern with a cleaner separation between arms: USP13 siRNA reduced SOX-2 to 0.47 of control at 48 h and 0.32 at 72 h, whereas alpelisib alone left it essentially unchanged (1.18 and 1.07), and the USP13 siRNA + alpelisib combination gave the lowest values recorded (0.41 at 48 h, 0.21 at 72 h). c-Myc levels declined in parallel over the same interval, with the weakest signal in the USP13-depleted and combination lanes at 48 and 72 h (Figure 5B). SK-MES-1 cells reproduced the qualitative pattern for all three proteins: progressive loss from 24 to 72 h, driven by the USP13 arm and most pronounced when alpelisib was added, although SOX-2 was expressed at markedly lower levels in this line than in NCI-H520 (Figure 5C).

Three findings in this data support the hypothesis as originally stated. First, the effect on all three effectors tracks the USP13 arm rather than the PI3Kα arm, which is the assignment the mechanism predicts: alpelisib blocks replenishment but cannot itself destabilize an existing protein pool. Second, a stability-driven mechanism would take 48-72 hrs to demonstrate an effect. Depletion becomes visible only as the pre-existing pool turns over, and it explains why the 48 h viability readout in Figure 4 captures the interaction at an early point in its development. Third, the combination lanes were consistently the lowest in the matrix without being dramatically lower than USP13 depletion alone, which matches the additive rather than strongly synergistic viability interaction and supports the framing of this pairing as one that reopens a therapeutic window for PI3Kα inhibition rather than one that produces supra-additive killing in its own right.

## 4 Discussion

### 4.1 Principal findings

EMET’s Agentic Research Director identified two mechanistically novel combination hypotheses for LUSC, each consistent with independent evidence validation, internal scientific review, and initial in vitro testing. Dual CDC7/PKMYT1 inhibition gave statistically significant synergy in both LUSC lines; dual USP13/PI3K inhibition gave a moderate but highly reproducible positive interaction in NCI-H520, with its predicted MCL-1/MYC/SOX2 mechanism validated. This is proof-of-concept for unbiased computational reasoning on an AI research environment: with no pre-specified target list, the system nominated untested pairs whose predicted mechanisms supported experimentally.

Two features of the CDC7/PKMYT1 data strengthen the mechanistic reading. First, the interaction grew from day 5 to day 7 in both lines, most markedly in NCI-H520, as expected of an effect compounding across cell cycles rather than acute cytotoxicity [37]. Second, synergy emerged unoptimized. Neither line was genotyped for CCNE1 amplification or RB1 loss, the strongest reported predictors of PKMYT1-pathway sensitivity [27, 38]. Hence a biomarker-selected model might show stronger synergy. In addition, an intervention strategy of CDC7 before PKMYT1 can also amplify the inhibitory effect. Finally, the growth inhibition observed in these assays warrants further resolution of the relative contributions of mitotic catastrophe versus a broader antiproliferative effect. A complementary panel comprising cell-cycle profiling, mitotic scoring, and micronuclei and γH2AX quantification would supply direct evidence for the proposed mechanism [37].

Mechanistic validation of the USP13/PI3K combination was confirmed by immunoblotting, which showed depletion of MCL-1, c-Myc and SOX-2, driven principally by the USP13 arm. USP13 is amplified in roughly half of LUSC tumours and sits within the 3q26 amplicon immediately adjacent to PIK3CA [33] and alongside SOX2; it stabilizes MYC by removing K48-linked ubiquitin, with USP13, MYC and SOX2 protein abundance positively correlated across human LUSC [35], and in other epithelial cancers it binds, deubiquitinates and stabilizes MCL-1 directly [36]. The therapeutic implication is specific to the PI3K inhibitor class, which has repeatedly failed as LUSC monotherapy [24], and the strategy has precedent: USP13 depletion sensitizes ovarian cancer cells to PI3K/AKT inhibition, where USP13 amplification has been proposed as a mechanism of intrinsic resistance to the class [33], and sensitizes cervical cancer cells to MCL-1-constrained BH3 mimetics [36]. USP13 is therefore a tractable route to lowering the survival threshold that PI3K inhibition alone does not reach. The additive rather than supra-additive interaction is what the rationale predicted, since the objective was to expand a therapeutic window rather than to produce synergistic killing. Quantifying that window will require evaluating alpelisib IC50 shifts under USP13 loss of function, target protein turnover, and stratification by 3q26 status, alongside an isoform-selective USP13 inhibitor to separate USP13 from the USP10 activity of spautin-1.

### 4.2 Why conventional mechanisms would have missed these combinations

Both pairings would have been hard to surface conventionally, for different reasons. Neither had been tested together in any LUSC line, placing them beyond supervised synergy prediction. Models trained on historical screens get no signal from an unprecedented pair. Nor would an unbiased screen prioritize them. CDC7 and PKMYT1 neither interact physically nor share a cell-cycle phase, and their convergence is visible only through CDK1 regulation, and only once TP53 loss has removed the G1 checkpoint. The pairing follows from checkpoint-architecture reasoning, not from pairwise similarity a screen or compound-centric heuristic would surface. USP13/PI3K is harder still: the predicted benefit is not supra-additive killing but reopening a therapeutic window for a class already failed in this histology - an objective a synergy score cannot detect and a synergy-ranked screen would actively discard. Reaching either manually would require one reviewer to weigh LUSC’s complex genetic landscape, checkpoint biology, deubiquitinase substrate specificity, and the PI3K clinical failure record against one another in a single judgement. The Director performs that conjunction systematically, and is correspondingly better at surfacing opportunities of this kind. Breadth of evidence is a second factor: with multi-omic, functional, pharmacological, clinical, and literature-derived data in one environment, support could be assembled across modalities normally queried separately, so a hypothesis weakly supported in any single modality could still be recognised as well supported overall.

### 4.3 Implications for combinational therapies

These results are best interpreted against how difficult combination identification remains by conventional means. The search space is the first obstacle: a 1,000-compound library already implies some 500,000 pairwise combinations before dosing ratios and scheduling are considered, placing exhaustive screening beyond practical and economic reach [39–41]; high-throughput platforms now test thousands of pairs routinely [42], but hit rates are low and the yield stays modest even among already-approved agents [43], against development economics in which around 90% of clinical programmes fail at over $2 billion per approval [2, 4]. The deeper limitation, however, is not throughput but explanation. Empirical screening establishes that a pair acts together without establishing why, and many combinations now in clinical use were arrived at empirically, their molecular basis reconstructed only in retrospect [41]. A screening score is a statement about one cell line under one set of conditions; a mechanistic account specifies which genotype should respond, which normal tissue is at risk, which schedule to use, what biomarker should stratify a trial, and what a negative result would mean. Synergy, the standard preclinical benchmark, is itself model-dependent and an imperfect proxy for clinical benefit [44], and several of the most effective approved combinations show none at all, benefiting patients through independent drug action instead [45, 46]. Combinations that arrive without such an account translate poorly, because nothing licenses the inference that an in vitro effect will survive into the clinic.

This is the gap the EMET platform is designed to close. Rather than scoring pre-specified pairs, it reasons across a multi-domain knowledge graph to generate rationalized, testable hypotheses prior to experimental testing. Both hypotheses here were supported by a mechanism: replication stress met with checkpoint loss in one case, and the removal of post-translational protein stabilization from a signaling axis that independently replenishes the same effectors in the other. That prior reasoning is what makes the two outcomes interpretable rather than merely positive or negative.

### 4.4 Broader applicability: repurposing, indication expansion, and the toolkits required

The need for such approaches is not confined to LUSC. Thousands of conditions still lack adequate therapy [1], and LUSC is a representative rather than exceptional case: a disease of substantial incidence in which decades of targeted-therapy trials have not reproduced the survival gains achieved in adenocarcinoma, leaving immunotherapy and chemotherapy as the backbone of systemic treatment [24]. For diseases defined by redundant, interconnected biology rather than a single dominant driver, combinations are not an optimization of monotherapy but a precondition for efficacy, and identifying them is precisely where conventional methods are weakest.

That backbone is itself instructive. Most agents that improved outcomes in squamous disease were developed for NSCLC broadly and benefit LUSC patients as a subgroup: consolidation durvalumab after chemoradiotherapy in unresectable stage III disease [47], adjuvant atezolizumab and adjuvant pembrolizumab following resection [48, 49], pembrolizumab with chemotherapy irrespective of PD-L1 expression [50], and ramucirumab in the second line [51]. Trials conducted specifically in squamous histology point the same way: nivolumab raised two-year survival from 8% to 23% over docetaxel [52], and pembrolizumab added to platinum–taxane chemotherapy extended median overall survival from 11.3 to 15.9 months [53]. The squamous-specific agents directed at a signalling axis rather than the immune system produced the smallest gains: necitmumab, an EGFR antibody, added 1.6 months of median overall survival [54], and afatinib, a pan-ErbB inhibitor, 1.1 months over erlotinib [55]. The Lung-MAP master protocol [21] tested this directly, assigning previously treated squamous patients to targeted agents against PIK3CA, FGFR, MET, cell-cycle and DNA-repair alterations; the targeted arms returned an overall response rate of 7% and a median overall survival of 5.9 months, against 7.7 months for docetaxel and 10.8 months for the immunotherapy arms [56]. The pattern is one of incremental, largely immune-mediated gains, with the disease’s own genomic architecture still substantially unexploited.

The same reasoning substrate extends naturally to two adjacent problems. Drug repurposing is structurally the same query run in the opposite direction: instead of asking which targets a disease exposes, one asks which diseases an existing agent’s mechanism should address, a question the knowledge graph already encodes, and the setting in which graph-based methods were first shown to work at scale [7]. Indication expansion is the narrower case of asking where an agent with established MoA, safety and pharmacology should be tried next, which is attractive precisely because it inherits a known clinical profile and so shortens the path from hypothesis to trial. Both benefit from the property that matters most in the present work: a candidate that arrives with an explicit mechanism specifies which patients should respond, and therefore how a trial should be stratified.

Realizing this requires integrating capabilities to extend the current system. Three are most consequential. First, patient-level and real-world evidence, e.g., trial outcomes, electronic health records, and pharmacovigilance signals, would let the Director reason about efficacy and safety in defined populations rather than in cell lines, and would let repurposing hypotheses be checked against observed outcomes in patients already taking the drug. Second, quantitative pharmacology, e.g., exposure–response modelling, tissue distribution, and drug–drug interaction prediction, is needed to convert a target-level hypothesis into a dosable regimen; the unresolved sequencing question raised in Section 4.1, and the additive myelosuppression risk a CDC7i + PKMYT1i regimen would be expected to carry, are exactly the class of problem this addresses, and neither can be settled from mechanism alone. Third, closing the loop experimentally: the Director already records outcomes in persistent memory, but its value compounds only if validation results at scale, including negative results, flow back as training and prioritization signal. Combining these with the reasoning architecture described here is what would turn a system that generates interpretable hypotheses into one that also learns which of its own mechanistic arguments prove reliable.

The results here are early and validated in two LUSC cell lines, and establish feasibility rather than clinical promise. Inference at scale, when it delivers mechanism alongside prediction, can nominate combination hypotheses that are novel, testable and interpretable, redirecting effort toward therapeutic approaches most likely to reach patients across the many conditions that still lack adequate therapy [1].

## 5 Supplementary Material

Supplementary Methods, Supplementary Results and Supplementary Figures 1–4 are provided in a separate Supplementary Material file accompanying this preprint.

