## supplementary materials for "Novel Target Combinations in Lung Squamous Cell Carcinoma proposed by the EMET AI Research Environment and supported by discovery stage experimentation"

*BenchSci*

### 1 Supplementary Methods

##### *1.0.1 Scheduling sequence of nodes in the graph-of-thoughts.*

The node cycle describes what happens within one hypothesis; it does not determine which hypothesis is worked on next. That scheduling decision is made by an outer exploration loop, summarized in Supplementary Figure 1, which alternates between distinct LLM roles and deterministic bookkeeping steps.

A round proceeds as follows. A *planner* inspects the current state of the graph and spawns new questions, expanding the set of nodes eligible for work — the frontier. A deterministic step then ranks that frontier by a branch score, so the order in which nodes are visited is fixed by an explicit, inspectable rule rather than by a model's discretion. *Research nodes* run in parallel over the highest-ranked entries on the frontier, each gathering evidence for its own assigned question and self-scoring what it finds. A second deterministic step aggregates those child scores upward to their parent nodes, so that a branch's value reflects what its descendants actually found rather than what it looked like when it was created; a line of reasoning that appeared promising but yielded weak evidence loses standing automatically. A *critic* then reviews the resulting material for quality and recommends which nodes to keep and which to drop — a judgement deliberately separated from the research nodes that produced and self-scored the evidence, so that assessment is not performed by the same process that has an interest in the outcome.

The round then writes to global memory before it ends. This is the step that makes the loop cumulative rather than merely iterative: the strategies attempted, the scores they attracted, the critic's keep-or-drop decisions, and any hypotheses set aside as already-known art are committed to a persistent store that outlives the round. Because the write happens inside the loop rather than only at the end of a mission, a later iteration can consult what earlier iterations established — which reasoning angles have already been exhausted, which regions of the graph have been sampled without yielding support, and which candidates were rejected and on what grounds. Two practical consequences follow. Effort is not spent re-deriving conclusions the system has already reached, since a question equivalent to one already answered can be recognized as such and pruned before research nodes are spawned against it. And the record is auditable after the fact: because outcomes are written as they occur rather than reconstructed at the end, the trace shows not only which hypotheses survived but which were considered and discarded, and why. The same store is what carries learning across whole rounds of a research mission, described in

Section 2.2.6.

Finally, a deterministic termination check ends the round if the frontier is empty or the iteration budget is exhausted; otherwise control returns to the planner and the graph is expanded again. When the loop does terminate, a *synthesizer* assembles the accumulated evidence into ranked hypotheses, which then pass to the validation and novelty filters described below.

Two features of this arrangement are worth drawing out. First, the division of labour between models and code is deliberate: every step that decides *what to explore next*, *how scores combine*, *what is committed to memory*, or *when to stop* is deterministic, while the LLM-driven stages are confined to generating questions, gathering evidence, and judging quality. Scheduling, bookkeeping, and termination are therefore reproducible and auditable, and the search cannot run away with itself because the iteration budget is enforced outside the model. Second, the stages are separated rather than collapsed into a single agent, which keeps planning, evidence gathering, and evaluation from being performed by one process in one pass.

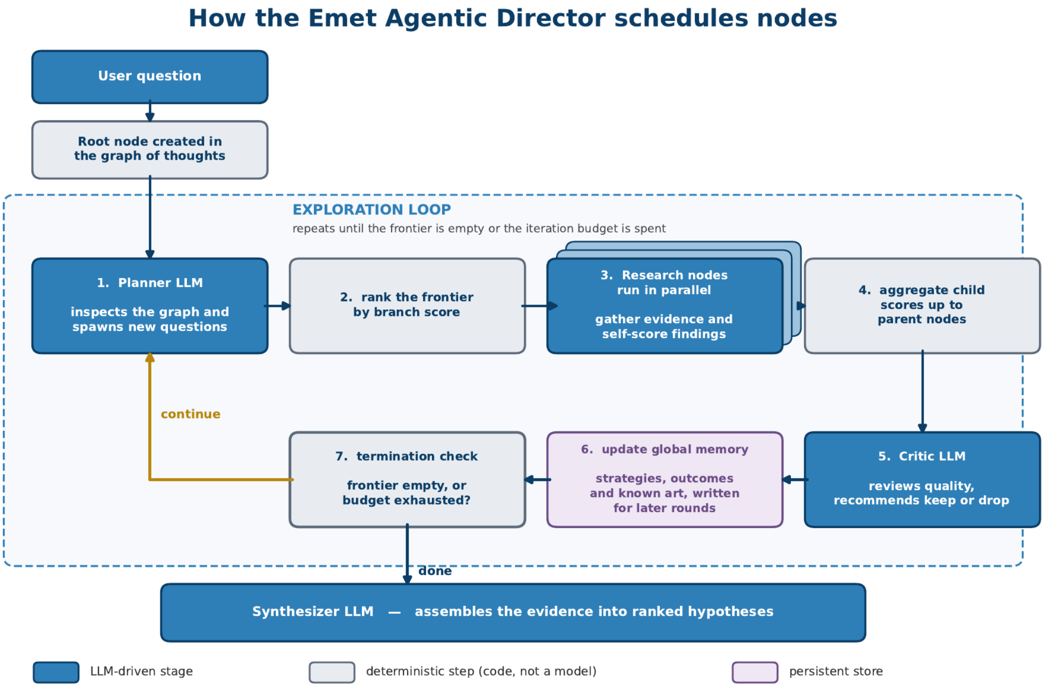


**Figure S1.** How the EMET Agentic Director schedules nodes. An outer exploration loop alternates between LLM-driven stages (blue) and deterministic steps implemented in code (grey). A planner spawns new questions onto the frontier (1); a deterministic step ranks that frontier by branch score (2); research nodes run in parallel, each gathering evidence for its own question and self-scoring what it finds (3); a second deterministic step aggregates child scores up to parent nodes, so branch standing reflects what descendants actually found (4); a critic then recommends which nodes to keep or drop (5). The round's strategies, scores, decisions and known art are committed to global memory (6, purple) before a deterministic termination check (7) either returns control to the planner or ends the round when the frontier is empty or the iteration budget is exhausted. Writing to memory inside the loop rather than only at its end is what lets later iterations avoid re-deriving established conclusions, and leaves an auditable record of what was discarded as well as what survived. Note that every decision about what to explore next, how scores combine, what is committed to memory, and when to stop is deterministic rather than delegated to a model.

### 2 Supplementary Results

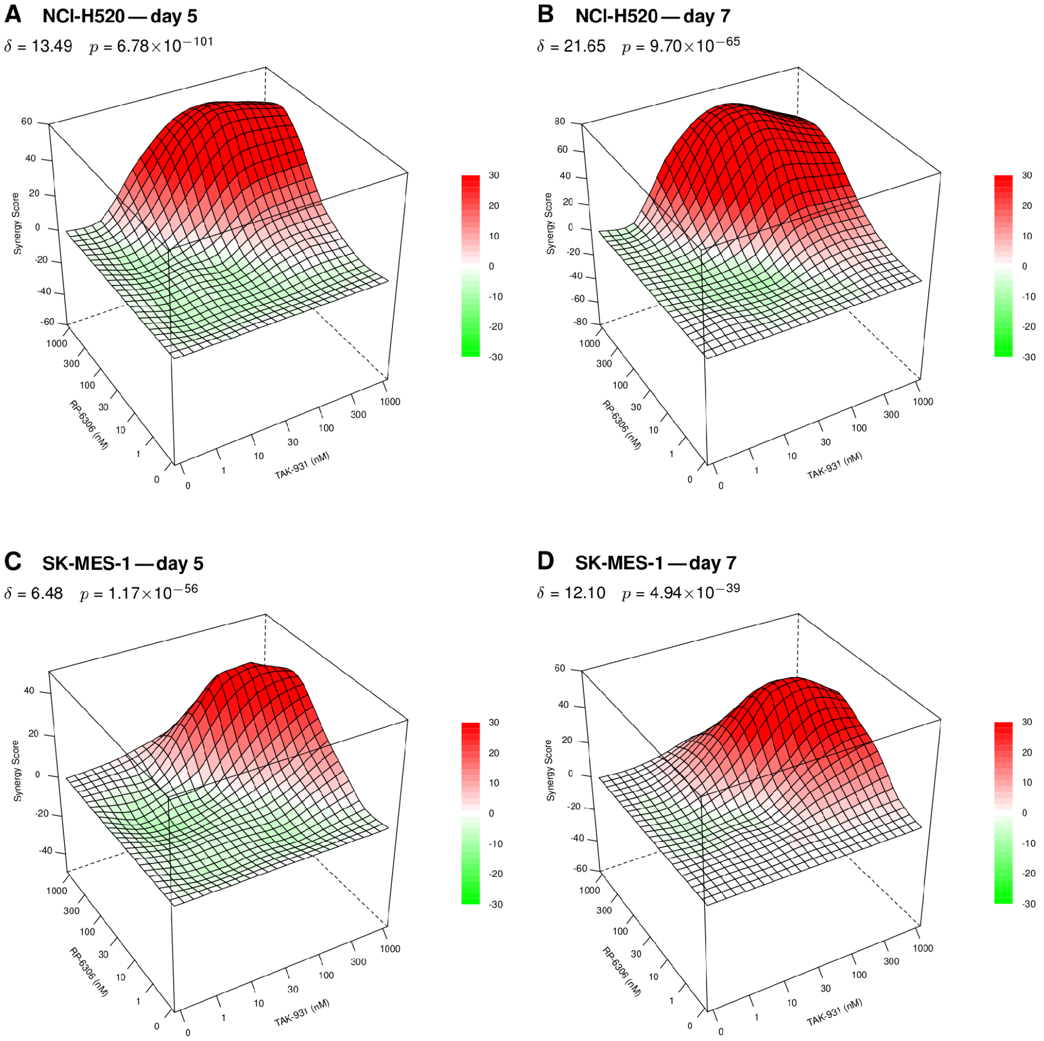


**Figure S2.** Bliss independence analysis reproduces the synergy and the day 5 to day 7 strengthening observed for CDC7 and PKMYT1 co-inhibition. Three-dimensional Bliss synergy score surfaces over the $7 \times 7$ TAK-931 × RP-6306 dose matrix (0, 1, 10, 30, 100, 300 and 1000 nM of each agent), recomputed in SynergyFinder+ [29, 31] under the Bliss independence reference model from the same 384-well CellTiter-Glo 2.0 viability data analysed under the HSA reference model in Figure 3. (A) NCI-H520 at day 5. (B) NCI-H520 at day 7. (C) SK-MES-1 at day 5. (D) SK-MES-1 at day 7. Above each surface, $\delta$ is the whole-matrix mean Bliss synergy score and p is the corresponding SynergyFinder+ p-value against a null of no interaction. The x and y axes give TAK-931 and RP-6306 concentration in nM and the z axis the Bliss synergy score at that dose pair; surfaces are coloured from green (antagonism) through white (no interaction) to red (synergy). By the thresholds applied in this study, scores below $-10$ indicate antagonism, scores between $-10$ and 10 an additive interaction, and scores above 10 synergy. Bliss places NCI-H520 in the synergistic range at both timepoints and agrees with HSA on the direction and time-dependence of the interaction in both lines; in SK-MES-1 at day 5 the Bliss score falls in the upper additive range (6.48) rather than the synergistic range returned by HSA (11.07), and crosses above 10 by day 7.

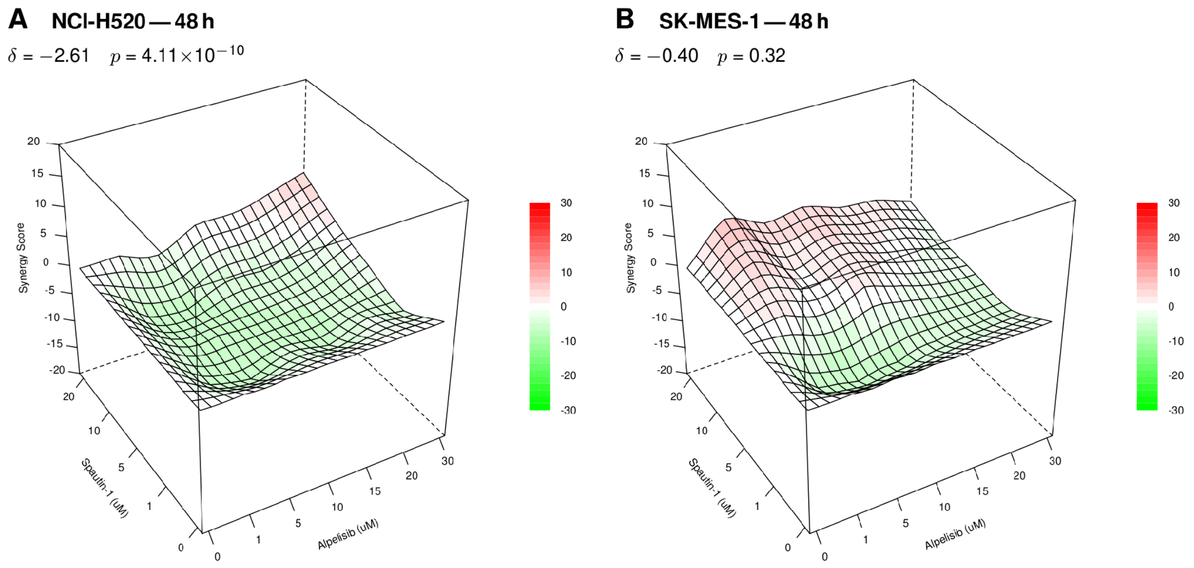


**Figure S3.** Bliss independence analysis places combined PI3Kα and USP13 inhibition close to independent drug action. Three-dimensional Bliss synergy score surfaces over the alpelisib × spautin-1 dose matrix (alpelisib 0, 1, 5, 10, 15, 20 and 30 μM; spautin-1 0, 1, 5, 10 and 20 μM) at 48 h, recomputed in SynergyFinder+ [29, 31] under the Bliss independence reference model from the same replicate-averaged 96-well CellTiter-Glo 2.0 viability data analysed under the HSA reference model in Figure 4. (A) NCI-H520. (B) SK-MES-1. Above each surface, δ is the whole-matrix mean Bliss synergy score and p is the corresponding SynergyFinder+ p-value against a null of no interaction. The x and y axes give alpelisib and spautin-1 concentration in μM and the z axis the Bliss synergy score at that dose pair; surfaces are coloured from green (antagonism) through white (no interaction) to red (synergy). Both surfaces sit close to zero over most of the matrix: the whole-matrix score is slightly negative in NCI-H520 (−2.61) and not distinguishable from zero in SK-MES-1 (−0.40, p = 0.32). Because both agents are individually active over much of this concentration range, Bliss sets a higher expected combination effect than the HSA model used in Figure 4, and the two models read together place this pairing close to independent drug action rather than synergy [45, 46].

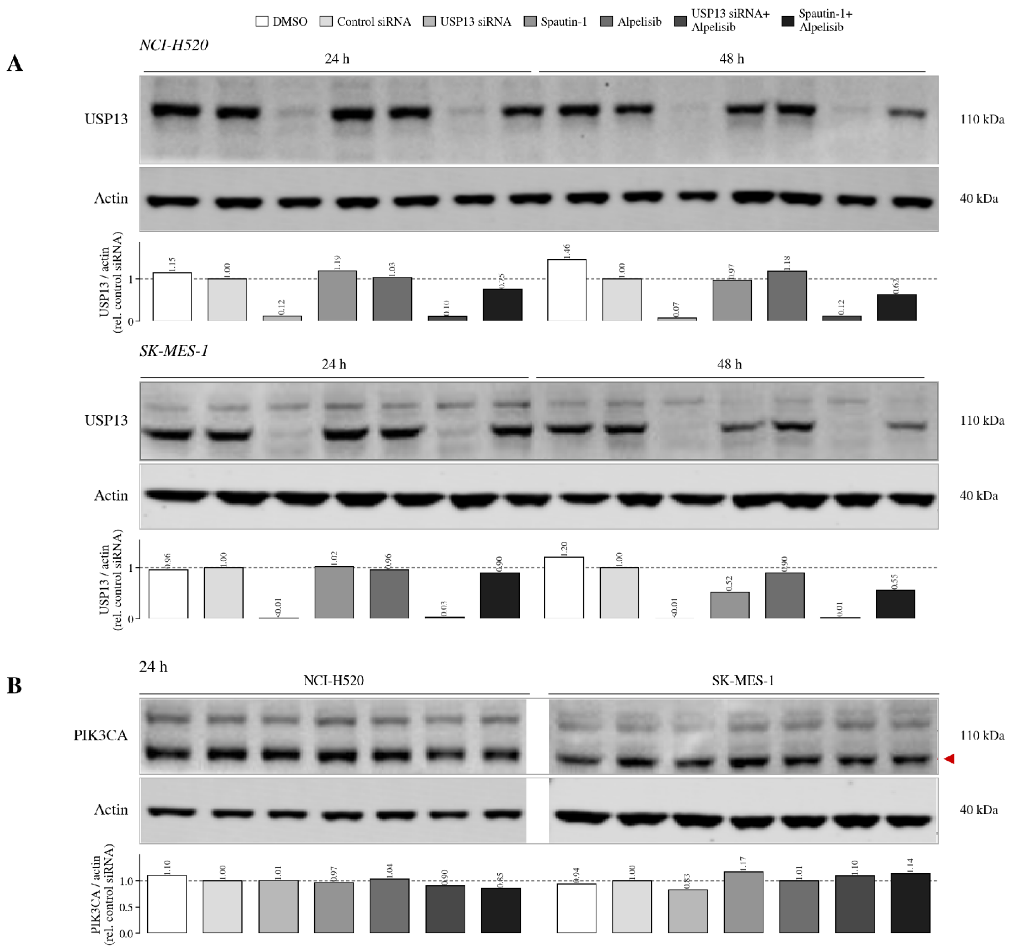


**Figure S4.** USP13 and PIK3CA protein levels across the USP13 loss-of-function and alpelisib treatment matrix in lung squamous cell carcinoma lines. Immunoblot analysis of the LUSC lines NCI-H520 and SK-MES-1 following USP13 depletion or inhibition, alone and in combination with the PI3Kα-selective inhibitor alpelisib. Within each block the seven lanes are, in order: (1) vehicle (DMSO); (2) non-targeting control siRNA; (3) USP13 siRNA; (4) 10 μM spautin-1; (5) 5 μM alpelisib; (6) USP13 siRNA + 5 μM alpelisib; (7) 10 μM spautin-1 + 5 μM alpelisib. Treatments are identified by the key above each blot. (A) USP13 (~110 kDa) in NCI-H520 (upper) and SK-MES-1 (lower) at 24 and 48 h. Both USP13 siRNA and spautin-1 reduce USP13 protein in each line, alone and together with alpelisib, establishing target engagement for the two approaches to USP13 loss of function used throughout this study. (B) PIK3CA (p110α, ~110 kDa) in NCI-H520 and SK-MES-1 under the same treatment matrix at 24 h. The two membrane images are separate exposures placed side by side and are delineated by the white gap between them. The red arrowhead marks the p110α band in SK-MES-1, which resolves close to a second immunoreactive species. Actin was probed on the corresponding membrane as the loading control for every panel, and approximate molecular weights are indicated to the right of each blot. Bar graphs beneath each panel show densitometric quantification normalized to actin and expressed relative to the non-targeting siRNA control (dashed line = 1.00). Primary antibodies are as listed in Table 2. This is the target-engagement evidence for the mechanistic analysis in Section 3.6; downstream effectors of this treatment matrix are shown in Figure 5.
